# Atomic-level evolutionary information improves protein-protein interface scoring

**DOI:** 10.1101/2020.10.26.355073

**Authors:** Chloé Quignot, Pierre Granger, Pablo Chacón, Raphael Guerois, Jessica Andreani

**Affiliations:** Université Paris-Saclay, CEA, CNRS, Institute for Integrative Biology of the Cell (I2BC), 91198, Gif-sur-Yvette, France; Department of Biological Chemical Physics, Rocasolano Institute of Physical Chemistry C.S.I.C, Madrid, Spain

**Keywords:** Protein-protein interactions, protein docking, protein scoring, protein evolution, protein structure, structural bioinformatics

## Abstract

The crucial role of protein interactions and the difficulty in characterising them experimentally strongly motivates the development of computational approaches for structural prediction. Even when protein-protein docking samples correct models, current scoring functions struggle to discriminate them from incorrect decoys. The previous incorporation of conservation and coevolution information has shown promise for improving protein-protein scoring. Here, we present a novel strategy to integrate atomic-level evolutionary information into different types of scoring functions to improve their docking discrimination.

We applied this general strategy to our residue-level statistical potential from InterEvScore and to two atomic-level scores, SOAP-PP and Rosetta interface score (ISC). Including evolutionary information from as few as ten homologous sequences improves the top 10 success rates of these individual scores by respectively 6.5, 6 and 13.5 percentage points, on a large benchmark of 752 docking cases. The best individual homology-enriched score reaches a top 10 success rate of 34.4%. A consensus approach based on the complementarity between different homology-enriched scores further increases the top 10 success rate to 40%.

All data used for benchmarking and scoring results, as well as pipelining scripts, are available at http://biodev.cea.fr/interevol/interevdata/

## 1 INTRODUCTION

Proteins are key actors in a great number of cellular functions and often work in collaboration with others, thereby forming interaction networks. Knowledge of the detailed 3D structure of protein-protein interfaces can help to better understand the mechanisms they are involved in. Difficulties in the experimental determination of protein assembly structures have prompted the development of *in silico* prediction strategies such as molecular docking. When no homologous interface structure can be identified and used as a template, free docking is used instead, involving a systematic search where many interface conformations called decoys are sampled (Huang, 2014; Porter, et al., 2019). These decoys are then scored according to properties such as interface physics, chemistry, and statistics (Huang, 2015; Moal, et al., 2013). Guided docking approaches integrating complementary sources of information are also becoming increasingly popular (Koukos and Bonvin, 2019).

Over time, protein interfaces are submitted to evolutionary pressure to maintain functional interactions. Thus, protein interfaces tend to be more conserved than other regions on their surface (Mintseris and Weng, 2005; Teichmann, 2002) and signs of coevolution can be detected at protein interfaces, where potentially disrupting mutations are compensated for with mutations in contacting positions on the protein partner. These phenomena of conservation and coevolution can provide useful information in the analysis and prediction of their 3D interface structures (Andreani, et al., 2020). For example, evolutionary information is at the heart of increasingly popular covariation-based approaches, such as statistical coupling analysis (SCA) (Socolich, et al., 2005) or direct coupling analysis (DCA) (Morcos, et al., 2011), for structural proximity prediction of residues based on multiple sequence alignments (MSAs). These approaches can be used to guide protein folding or to supplement predictions of macromolecular interactions (Cocco, et al., 2018; Cong, et al., 2019; Simkovic, et al., 2017). The vast majority of protein interaction site predictors successfully use evolutionary information, be it by sequence conservation, sequence co-evolution, or through homologous structures (Andreani, et al., 2020).

Evolutionary information can also be especially useful to guide molecular docking (Geng, et al., 2019). The InterEvDock2 server implements a docking pipeline that uses evolutionary information (Quignot, et al., 2018; Yu, et al., 2016). It takes advantage of the spherical Fourier-based rigid-body docking programme FRODOCK2.1 (Ramírez-Aportela, et al., 2016) for the sampling step and hands out a set of ten most probable interfaces based on a consensus between three different scores, FRODOCK2.1’s mostly physics-based score, SOAP-PP’s atomic statistical potential (Dong, et al., 2013) and InterEvScore (Andreani, et al., 2013). InterEvScore extracts co-evolutionary information from joint multiple sequence alignments of the binding partners (called coMSAs), but unlike covariation-based approaches such as DCA cited above, InterEvScore needs only a small number of homologous sequences to improve discrimination between correct and incorrect decoys, by combining coMSAs with a multi-body residue-level statistical potential. As seen in the benchmarking of InterEvDock2, InterEvScore presents a high complementarity with SOAP-PP (Quignot, et al., 2018). As both scores are based on statistical potentials but SOAP-PP has an atomic level of detail, we hypothesised that a score integrating evolutionary information at an atomic scale might pick up on finer properties to better distinguish near-natives from the rest of the decoys.

In InterEvScore, evolutionary information is given implicitly at residue-level through coMSAs and combined with a coarse-grained statistical potential. A major challenge in deriving evolutionary information to an atomic level of detail is finding a suitable way of representing residue-scale information from coMSAs at an atomic level. Here, we present a novel strategy to couple evolutionary information with atomic scores to improve decoy discrimination. We reconstruct an equivalent and hypothetical interfacial atomic contact network for each interface decoy and each pair of homologs present in the coMSAs, by using a threading-like strategy to generate explicit backbone and side-chain coordinates. These models can, in turn, be scored with non-evolutionary atomic-resolution scoring functions such as SOAP-PP (Dong, et al., 2013) or Rosetta interface score (ISC) (Chaudhury, et al., 2011; Gray, et al., 2003).

Here, we show that including explicit evolutionary information improves the top 10 success rate of SOAP-PP and ISC by 6 and 13 percentage points respectively, on a large benchmark of 752 docking cases for which evolutionary information can be used (Yu and Guerois, 2016). It also improves the top 10 success rate of the residue-level statistical potential from InterEvScore by 6.5 percentage points. We then use a consensus approach to take advantage of the complementarity between different scores. The top 10 success rate of a consensus integrating FRODOCK2.1 with InterEvScore and SOAP-PP increases from 32% to 36% when including the homology-enriched score variants. A more time-consuming consensus combining all scores with an explicit homolog representation reaches 40% top 10 success rate.

## 2 METHODS

### 2.1 Docking benchmark

We evaluated docking methods using the large docking benchmark PPI4DOCK (Yu and Guerois, 2016), where unbound structures unavailable from experiments were modelled by homology from unbound homologous templates. Each case in PPI4DOCK is associated to a coMSA, i.e. a pair of joint MSAs for the two docking partners. To focus on cases with enough co-evolutionary information, we excluded antigen-antibody interactions and cases with less than 10 sequences in their coMSAs. Sampling was performed using FRODOCK2.1 (see supplementary methods section 5.1.1) and only the top 10,000 decoys were kept. Near-native decoys were defined as being of Acceptable or better quality following the criteria from CAPRI (Critical Assessment of PRediction of Interactions) (Mendez, et al., 2003). To focus the study on scoring performance, only cases that have a near-native within the top 10,000 FRODOCK2.1 decoys were used for benchmarking. This resulted in a final benchmark of 752 cases (supplementary Table 5-1).

Performance was measured by top N success rate, i.e. the percentage of cases with at least one near-native in the top N ranked decoys. We especially focus on the top 10 success rate traditionally used as a docking metric, and the top 50 success rate since consensus computation typically involves the top 50 decoys of each score (see section 2.2.1). Additional metrics are available in the supplementary information (section 5.1.2).

### 2.2 Scoring functions

In addition to FRODOCK2.1’s integrated score (Ramírez-Aportela, et al., 2016), we rescored decoys and their threaded homologs with InterEvScore, SOAP-PP, and Rosetta interface score (ISC).

InterEvScore combines co-evolutionary information taken from coMSAs with a residue-level statistical potential (Andreani, et al., 2013). It was re-implemented to accelerate the scoring step (see supplementary methods section 5.1.3).

SOAP-PP is an atomic statistical-based score integrating distance-dependent potentials learnt on a set of real complex structures and normalised on a set of incorrect PatchDock decoys (Dong, et al., 2013). Here, we use a faster in-house implementation of this score (see supplementary methods section 5.1.3).

Rosetta interface score (ISC) includes a linear combination of non-bonded atom-pair interaction energies and empirical and statistical potentials among other terms (Chaudhury, et al., 2011; Gray, et al., 2003). This score is calculated by subtracting the total energy of both monomeric structures from the total energy of the complex structure. Since Rosetta ISC is sensitive to small variations and clashes at the interface, we included high-resolution interface side-chain optimisation as a scoring option (see supplementary methods section 5.1.3). Decoys for which Rosetta scoring did not converge after 10 iterations were assigned the worst score for that case. As Rosetta ISC scoring can take up to a couple of minutes per structure, we scored only the top 1,000 FRODOCK2.1 decoys (noted later 1k) per case rather than 10,000 (noted 10k).

#### 2.2.1 Consensus scores

The aim of the consensus is to preferentially select decoys supported by several scores. Consensus calculations were performed similarly to InterEvDock2 (Quignot, et al., 2018) to obtain a set of 10 most likely decoys depending on the agreement between several scoring functions. Here, we apply consensus scoring to combinations of 3 to 5 different scoring functions. For a given set of scoring functions, ordered according to their individual performances from best to worst performing, the top 10 decoys of each scoring function receive a convergence count based on the number of similar decoys (defined as L-RMSD ≤ 10 Å) that are found in the top 50 decoys of each other scoring function. The final 10 consensus decoys are selected iteratively by decreasing convergence count (if > 1). In the case of a tie, decoys are selected according to the ranking order of their respective scoring functions. Note that decoys are added to the top 10 consensus only if they are not structurally redundant with the already selected ones (L-RMSD > 10 Å). If necessary, the consensus list is completed up to 10 decoys by selecting the top 4, 3, 3 decoys for a consensus between three scoring functions (or the top 3, 3, 2, 2 or top 2, 2, 2, 2, 2 decoys for a consensus between four or five scoring functions, respectively).

### 2.3 Docking strategy to integrate evolutionary information

The proposed homology-enriched docking pipeline consists of four steps outlined in Figure 2-1. First, we dock query proteins A and B for which we are trying to predict the 3D structure of the complex using FRODOCK2.1 (Ramírez-Aportela, et al., 2016). This results in a set of rotational and translational transforms that define a maximum of 10,000 complex decoys (Figure 2-1A). In parallel, we construct coMSAs and subsample them to a subset of M pairs of homologs (proteins A_1_ and B_1_, A_2_ and B_2_, …, A_M_ and B_M_, homologs of query proteins A and B respectively) (see section 2.3.1). We model the unbound structures of this subset of M pairs of homologs, using the threading function from RosettaCM’s pipeline (Song, et al., 2013) and the unbound query protein structures as templates (see Figure 2-1B and section 2.3.2). We then generate complex equivalents to each query decoy by applying the translational and rotational transforms obtained in the docking step to each pair of homologs. Figure 2-1C illustrates this reconstruction for the first pair of homologs (proteins A_1_ and B_1_). Finally, we average scores over the query decoy itself and its equivalent homolog decoys to obtain a final per-decoy score that integrates all the information (Figure 2-1D). Note that for one case, we have to compute (M+1) x N scores to obtain the final ranking of N decoys. The scoring functions we used are described in section 2.2. All steps of the pipeline are easily parallelisable to reduce end-user runtime, whether through MPI (sampling step) or by splitting over decoys (scoring steps).

**Figure 2-1:**
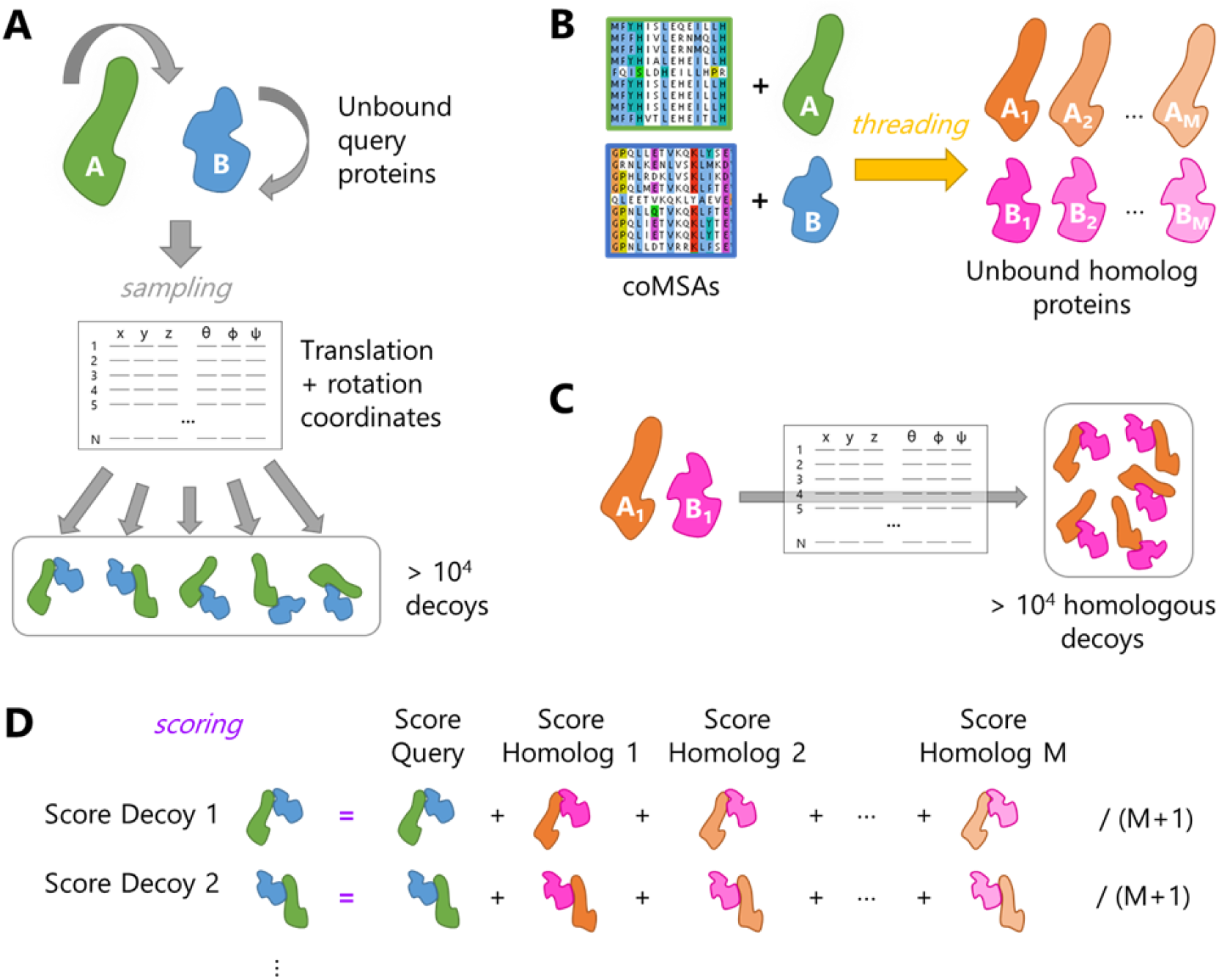
Docking pipeline with explicit modelling of decoy homologs. (A) Upon docking of query unbound structures (proteins A and B in green and blue), FRODOCK2.1 outputs a rotation and translation matrix to reconstruct the corresponding decoys. (B) To generate their homologous counterparts, the unbound structures of each homolog (proteins A_1_ and B_1_, A_2_ and B_2_, …, A_M_ and B_M_, in various shades of orange and magenta) are threaded based on the query unbound structures (proteins A and B) and the homologous sequence alignments in the coMSAs of the query proteins. (C) For each homolog pair (such as homolog 1 illustrated here), decoys can be reconstructed using the same rotation and translation matrix as for the query. (D) The final score of each decoy (left column) corresponds to the average score over itself and its M homolog equivalents for a given scoring function.

#### 2.3.1 Subsampling homologs in the coMSAs

Homologous sequences used in scoring were taken from the coMSAs provided with the PPI4DOCK benchmark, reduced to maximum M=40, and then to M=10 sequences (plus the query sequence) to limit computational time. Indeed, it was already seen with InterEvScore that co-evolutionary information can be extracted from alignments with as few as 10 sequences (Andreani, et al., 2013). The sequences in the coMSAs are ordered by decreasing average sequence identity with the query sequences. This is taken into account when sub-selecting sequences to keep a representative subset of sequences in both reduced coMSAs. Sequence selection was performed in three steps. First, the number of sequences was cut at 100, as in the InterEvDock2 pipeline. Then the alignment was filtered with hhfilter 3.0.3 (Remmert, et al., 2011) from the hh-suite package. hhfilter was applied with the “-diff X” option on the concatenated coMSAs and the value of X was adjusted for each case to return a reduced alignment with no more than 41 sequences (i.e. the query + 40 homologs). At this stage, we obtain the first set of reduced coMSAs with maximum 40 sequences, which we call coMSA^40^, and that are representative of the full diversity of the initial coMSAs. Finally, 11 equally distributed sequences (i.e. the query + 10 homologs) were uniformly selected within coMSA^40^ in order to preserve sequence diversity compared to the initial coMSAs (see supplementary methods section 5.1.4). The final set of reduced coMSAs is called coMSA^10^.

#### 2.3.2 Threading models

Pairwise alignments between the template structure and the homolog sequence to be modelled were directly extracted from the reduced coMSAs. The templates used for threading were the unbound template structures provided in the PPI4DOCK benchmark (Yu and Guerois, 2016) (see supplementary methods section 5.1.5).

Rosetta’s threading programme, the first step in the RosettaCM pipeline (Song, et al., 2013), was used to thread the homologous sequences onto the template structure. We used Rosetta 3.8 (version 2017.08.59291). No insertion, N- or C-terminus were modelled. This resulted in gapped and mainly structurally conserved threaded models of the homologs, where backbone coordinates remained unchanged and side-chain rotamers were different from the template’s side-chains only if the residue type changed between the template and the homologous sequence (Figure 2-1B).

## 3 RESULTS

### 3.1 Consensus approach with implicit homology scoring

In previous work, we integrated evolutionary information implicitly at the coarse-grained level by scoring decoys with residue-based InterEvScore (noted IES) (Andreani, et al., 2013). In IES, for each decoy, we enumerate all residue-level interface contacts. We then use a res-idue-level statistical potential to score decoys by considering all sequences in a coMSA and assuming the same contacts were conserved in all homologous interfaces.

We also combined InterEvScore with complementary scores FRODOCK2.1 and SOAP-PP (supplementary Figure 5-1A) in a three-way consensus score, denoted Cons^3^, which preferentially selects decoys supported by several scores (section 2.2.1) (Quignot, et al., 2018; Yu, et al., 2016). Compared to individual scores, we observed a notable boost of about 8 points in the top 10 success rate using Cons^3^, which captures a near-native in the top 10 decoys in 32% of the cases (Table 3-1 and Figure 3-1A).

**Table 3-1:** Performance of consensus scores including InterEvScore implicit homology scoring. Scores used in three-way consensus score Cons^3^ were SOAP-PP on the top 10,000 FRODOCK2.1 decoys (SPP/10k), InterEvScore on full coMSAs and on the top 10,000 FRODOCK2.1 decoys (IES/10k) and FRODOCK2.1 (FD2.1). Performances of individual scores used in the consensus are reported in terms of top 10 and top 50 success rates since consensus calculation relies on the top 50 decoys ranked by each component score.

| Score | Top 10 success rate | Top 50 success rate |
| --- | --- | --- |
| **FD2.1** | 164 (21.8%) | 292 (38.8%) |
| **IES/10k** | 182 (24.2%) | 287 (38.2%) |
| **SPP/10k** | 183 (24.3%) | 328 (43.6%) |
| **Cons<sup>3</sup>** | **241 (32.0%)** | / |

**Figure 3-1:**
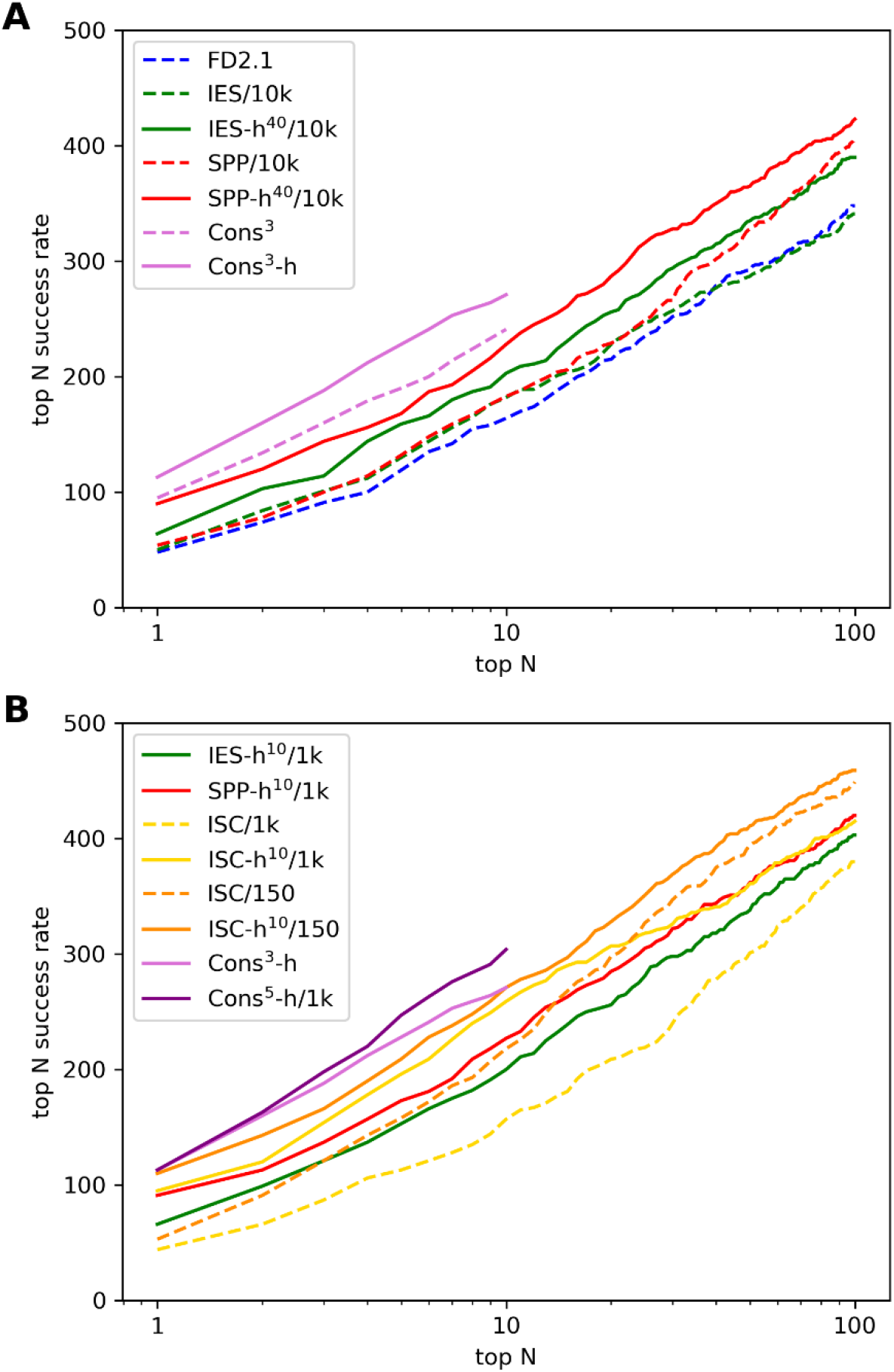
Success rate as a function of the number of selected decoys for individual and consensus scores. Illustration of the success rate on an increasing number of top N decoys with N going from 1 to 100. (A) FRODOCK2.1 (FD2.1), SOAP-PP (SPP) and InterEvScore (IES) individual and consensus scores (dashed lines) and their homology-enriched variants on coMSA^40^ and 10,000 decoys (10k) (solid lines). (B) Rosetta ISC scores (dashed lines) together with homology-enriched variants of individual scores on coMSA^10^ and 1,000 decoys (1k) and selected homology-enriched consensus scores (solid lines). Performances were measured on 752 benchmark cases. Note that consensus scores produce only a selection of 10 decoys, hence they stop at N=10.

This complementarity between the examined scores, in particular SOAP-PP and InterEvScore, (supplementary Figure 5-1A) prompted us to attempt a more explicit integration of evolutionary information into the various scores. Following the pipeline described in methods section 2.1 (Figure 2-1), in the next sections, we include evolutionary information into individual scores InterEvScore and SOAP-PP through explicit atomic-level models of homologous decoys.

### 3.2 InterEvScore with explicitly modelled homologs

For efficiency, we represent homologs at atomic resolution by threading their sequences onto the query structure (section 2.3.2). As a first step to validate this new representation of evolutionary information, we test the performance of InterEvScore on these threaded models and compare it with the original InterEvScore. With the threaded models, atomic contacts are re-defined in each homolog at an explicit level, rather than implicitly deduced from the coMSAs as in the original InterEvScore. In practice, we calculate the threaded homolog version of InterEvScore (denoted IES-h) by scoring query decoys and their threaded homolog equivalents with the InterEvScore statistical potentials (section 2.3). The final score of each query decoy corresponds to the average score over the query decoy itself and its homologs.

Table 3-2 and Figure 3-1A show the performance of IES-h^40^, i.e. IES-h computed using threaded homologs from the set of reduced coMSAs with a maximum of 40 sequences (coMSA^40^, see section 2.3.1). Results for the original InterEvScore with complete coMSAs (IES) and coMSAs^40^ (IES^40^) are also shown for comparison. Reducing the number of sequences to 40 does not strongly affect performance in terms of the top 10 and top 50 success rates. However, the top 10 success rate increases from 23.8% to 27.0% when using explicit threaded models (IES-h^40^) instead of only implicit coMSA information (IES^40^). Of note, a variant of InterEvScore without evolutionary information, where only the query decoy gets scored by the statistical potential, has a much lower top 10 success rate of 20.5% (supplementary Table 5-2).

**Table 3-2:** Performance of InterEvScore using coMSAs without or with threaded models. Top 10 and top 50 success rates of InterEvScore on complete coMSAs (IES, reported in section 3.1 and Table 3-1) and coMSA^40^ (IES^40^) compared to InterEvScore using explicit threaded models of homologs in coMSA^40^ (IES-h^40^) on 10,000 decoys (/10k). Performances were measured on 752 benchmark cases.

|  | Top 10 success rate | Top 50 success rate |
| --- | --- | --- |
| **IES/10k** | 182 (24.2%) | 287 (38.2%) |
| **IES<sup>40</sup>/10k** | 179 (23.8%) | 284 (37.8%) |
| **IES-h<sup>40</sup>/10k** | **203 (27.0%)** | **335 (44.5%)** |

The difference in performance between IES^40^/10k and IES-h^40^/10k can be explained by the fact that, in IES-h^40^, contacts are not extrapolated from the query interface network anymore but are redefined in each homolog based on their modelled interface structure.

### 3.3 Homology-enriched SOAP-PP

Having explicit structures at atomic resolution corresponding to each homolog enables us to score them directly using an atomic potential such as SOAP-PP (Dong, et al., 2013), which might be able to better exploit the atomic detail of homologs for the final ranking of query decoys. As for the threaded version of InterEvScore, homology-enriched SOAP-PP (SPP-h^40^) consists in the average SOAP-PP score over all homologs including the query decoy itself.

SPP-h^40^ performs better than SOAP-PP on the query decoys alone (Table 3-3 and Figure 3-1A). Using threaded homology models in this way gives a large performance boost to SOAP-PP (+6 percentage points on the top 10 success rate). SPP-h^40^ also outperforms InterEvScore and IES-h^40^ (section 3.2) as well as the FRODOCK2.1 score (section 3.1).

**Table 3-3:** Performance of SOAP-PP against SPP-h^40^. Top 10 and top 50 success rates of SOAP-PP (SPP) compared to its homology-enriched version SPP-h^40^ over sequences in coMSA^40^ on 10,000 decoys (/10k). Performances were measured on 752 benchmark cases.

|  | Top 10 success rate | Top 50 success rate |
| --- | --- | --- |
| **SPP/10k** | 183 (24.3%) | 328 (43.6%) |
| **SPP-h<sup>40</sup>/10k** | **228 (30.3%)** | **365 (48.5%)** |

### 3.4 Homology-enriched Rosetta interface score (ISC)

Since we build atomic-level homolog models of decoys, we can score them explicitly using a physics-based score such as Rosetta ISC. As Rosetta scoring is much more computationally expensive (about 750 times slower) than SOAP-PP and InterEvScore, to compute homology-enriched ISC, the number of decoys was reduced to 1,000 (as ranked by FRODOCK2.1) and the number of homologs to 10 (coMSAs^10^, section 2.3.1).

As above, homology-enriched ISC consisted in the average score of the query and its homologous decoys (ISC-h^10^). For easier comparison, homology-enriched InterEvScore and SOAP-PP were evaluated in the same conditions (*i.e.* 1,000 decoys and coMSAs^10^) (Table 3-4 and Figure 3-1B). Their success rates are very similar to those with 10,000 decoys and coMSAs^40^ (supplementary Table 5-3). Even though ISC on query decoys performs worse than SPP-h and IES-h, ISC-h^10^ largely outperforms the best-performing individual score, SPP-h^10^, with 34.4% top 10 success rate (259 cases) compared to 30.2% (227). With only 165 successful cases in common, SPP-h^10^ and ISC-h^10^ remain very complementary (supplementary Figure 5-1B).

**Table 3-4:**
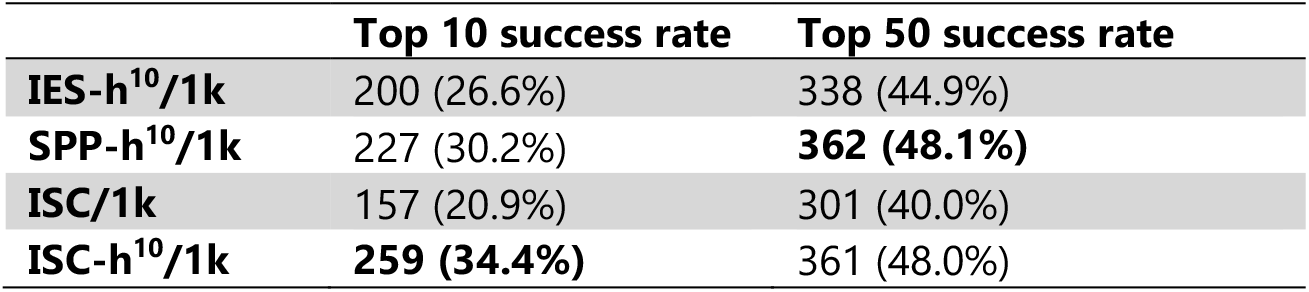
Scoring performance of Rosetta homology-enriched ISC. Scoring performance of ISC on query decoys only and using the threaded homology models (ISC-h^10^) on top 1,000 FRODOCK2.1 decoys (1k) and coMSA^10^ as well as the performance of SPP-h^10^ and IES-h^10^ on 1,000 FRODOCK2.1 decoys with coMSA^10^ for easier comparison. Performances were measured as the top 10 and top 50 success rates on 752 benchmark cases.

Note that for scores calculated on the top 1,000 FRODOCK2.1 decoys, success rates are technically capped to 77.1%, as only 580 cases out of the 752 in our benchmark have a near-native within this subset of decoys. In light of this, the ISC-h^10^/1k performance is all the more remarkable.

#### 3.4.1 Using ISC to re-score homology-enriched decoys

ISC-h^10^ showed the highest top 10 success rate from all scores tested above, but scoring 1,000 × 11 decoys with Rosetta ISC is excessively time consuming in a generalised docking context as it takes approximatively 137 CPU hours per case (supplementary Table 5-4). One way to alleviate the total scoring time is to score only a pre-selected amount of decoys, using Rosetta ISC as a second step in the scoring pipeline.

In Cons^3^, we pre-selected the top 50 decoys of FRODOCK2.1, InterEvScore, and SOAP-PP. Similarly, here we use the top 50 decoys of the top-performing homology-enriched score variants tested above, namely SPP-h^40^/10k and IES-h^40^/10k, as well as FRODOCK2.1. These scores have a high complementarity in terms of top 10 success rate with only 67 cases found in common between all three (supplementary Figure 5-1C). Using this subset of 150 pre-selected decoys for ISC scoring (referred to with /150h) reduced scoring times approximately by a factor 7. We enrich near-natives in this set of 150 decoys since they were pre-selected by three already well-performing scores, but only 476 out of 752 cases in our benchmark possess a near-native in this subset.

In terms of the top 10 success rate, both ISC-h^10^ and ISC perform better on 150 than 1,000 decoys with 36.0% and 29.0% top 10 success rate instead of 34.4% and 20.9%, respectively (Table 3-5 and Figure 3-1B). Here again, the addition of evolutionary information to ISC through the threaded homolog models remarkably increases its performance. ISC-h^10^/150h has the best performance of all tested scores so far, for a much lower computational cost than ISC-h^10^/1k.

**Table 3-5:** Performance of ISC and ISC-h^10^ on 150 pre-selected decoys. Below are summarised the top 10 success rates of ISC and ISC-h^10^. Top 10 success rates of ISC/150h and ISC-h^10^/150h were calculated after a pre-selection of a maximum of 150 decoys taken from the 3 x top 50 decoys of IES-h^40^/10k, SPP-h^40^/10k, and FRODOCK2.1. Scoring was performed on all 752 benchmark cases.

| Score | Top 10 success rate | Top 50 success rate |
| --- | --- | --- |
| ISC/150h | 218 (29.0%) | 394 (52.4%) |
| ISC-h<sup>10</sup>/150h | 271 (36.0%) | 411 (54.7%) |

### 3.5 Homology-enriched consensus scoring

As a first step, we calculate Cons^3^-h, the homology-enriched variant of the Cons^3^ base consensus score presented in section 3.1. Calculating a three-way consensus using higher-performing homology-enriched variants (Cons^3^-h) instead of their original counterparts (Cons^3^) increases the top 10 success rate from 32% to 36% (Table 3-6 and Figure 3-1A). Consensus Cons^3^-h performs as well as ISC-h^10^/150h, while calculated on the same top 150 decoys, and computation is about 20 times faster for Cons^3^-h than for ISC-h^10^/150h.

**Table 3-6:** Performance of homology-enriched consensus scores. Performance of three-, four- and five-way consensus scores in terms of top 10 success rates on 752 benchmark cases and approximate timescales for the whole pipeline (including sampling with FRODOCK2.1, homology model generation, scoring steps, and consensus calculation). Scores used in Cons^3^ were SOAP-PP/10k, InterEvScore/10k, and FRODOCK2.1. Scores used in all homology-based consensuses (Cons^X^-h) were FRODOCK2.1, SPP-h^40^/10k, IES-h^40^/10k, ISC and ISC-h^10^. The three-way consensus included the first three scores, four-way consensuses included all scores up to ISC and five-way consensuses included all of them. Cons^X^-h/150h included ISC scores over 150 decoys only and Cons^X^-h/1k included ISC scores over 1k decoys.

| Consensus | Top 10 success rate | Whole pipeline time estimates on 4 CPU* |
| --- | --- | --- |
| Cons<sup>3</sup> | 241 (32.0%) | 15 min |
| Cons<sup>3</sup>-h | 271 (36.0%) | 15 min |
| Cons<sup>4</sup>-h/150h | 276 (36.7%) | 45 min |
| Cons<sup>4</sup>-h/1k | 282 (37.5%) | 3 h 15 |
| Cons<sup>5</sup>-h/150h | 289 (38.4%) | 5 h 30 |
| Cons<sup>5</sup>-h/1k | 304 (40.4%) | 34 h 30 |

\* all steps are parallelisable using MPI (sampling) or over the decoys (scoring)

Out of the 271 successful cases for Cons^3^-h and ISC-h^10^/150h, only 199 cases are common. Moreover, ISC and ISC-h^10^ remain complementary to SPP-h^40^/10k, IES-h^40^/10k, and FRO-DOCK2.1 (supplementary Figure 5-1D and Figure 5-1E). This led us to test four- and five-way consensus approaches to combine ISC optimally with other homology-enriched scores. We tested two four-way consensuses that integrate ISC without homology on 1,000 or 150 de-coys (Cons^4^-h/1k and Cons^4^-h/150h respectively) and two five-way consensuses that inte-grate ISC both with and without homology on 1,000 or 150 decoys (Cons^5^-h/1k and Cons^5^-h/150h respectively). Performances are reported in Figure 3-1B and Table 3-6, together with time estimates when parallelising the whole pipeline on 4 CPUs.

With five-way consensus Cons^5^-h/1k, the top 10 success rate rises to 304 cases (40.4%). Unfortunately, computation time strongly increases, since we have to compute ISC-h^10^ on 1,000 decoys. The most time-effective consensus, Cons^3^-h, has 36.0% top 10 success rate and the same top 1 success rate as Cons^5^-h/1k (Figure 3-1B and supplementary Table 5-5).

## 4 DISCUSSION

In InterEvScore (Andreani, et al., 2013), evolutionary information improved protein-protein scoring performance when given implicitly through coMSAs and coupled with a coarse-grained, residue-level statistical potential. Combining InterEvScore with complementary scoring functions FRODOCK2.1 and SOAP-PP by computing a consensus (Quignot, et al., 2018; Yu, et al., 2016) improved over the individual scores, reaching 32% top 10 success rate (see Table 3-1). However, this strategy did not take full advantage of the three scores’ complementarity and we thus decided to combine directly evolutionary information from coMSAs with atomic scores such as SOAP-PP. To this aim, we threaded coMSA homologs of docked query proteins and scored homologous decoys together with each query decoy.

With this new explicit implementation of evolutionary information, we tested a variant of InterEvScore where we scored decoys and their modelled homologs with a residue-level statistical potential. This modified version (named IES-h) had a slightly improved success rate compared to the implicit homology version (see Table 3-2). The explicit representation of homologous decoys enabled us to build homology-enriched versions of atomic scores SOAP-PP (SPP-h) and Rosetta ISC (ISC-h). For both, adding homology drastically improved top 10 success rates (see Table 3-3 and Table 3-4) even when coMSAs were down-sampled to a maximum of 10 homologous sequences. The Rosetta homology-enriched version, ISC-h^10^, had outstanding performances, but it also was the most time-consuming score, about 750 times slower than SOAP-PP or InterEvScore. The first compromise between computation time and performance was to run ISC-h^10^ on a pre-selection of 150 decoys defined by the top 50 decoys of SPP-h^40^/10k, IES-h^40^/10k, and FRODOCK2.1 (see Table 3-5). This score had the same top 10 success rate (36%) as a much faster consensus score involving the same top 150 decoys. Taking further advantage of this complementarity, different four- and five-way consensus calculations managed top 10 success rates from 36.7% to 40.4% at runtimes ranging from 45 minutes to 34.5 hours on four CPUs (Table 3-6).

Our homology enriched scoring scheme is robust to change in the definition of near-natives (supplementary Table 5-6) and in evaluation metrics (supplementary Table 5-7). Using a more stringent definition of near-natives (as being of at least Medium quality according to CAPRI criteria) still allows homology enrichment to boost predictive performance of scoring functions. However, consensus scores become less efficient than the best individual scoring functions, probably because when grouping decoys with a relatively loose similarity criterion (see methods section 2.2.1), we do not manage to selectively up-rank Medium quality decoys (supplementary Table 5-6).

We further tried to understand the origin of the large performance improvements obtained through homology enrichment. Scoring performance improves when near-natives are recognised better (positive selection) or when wrong decoys are down-ranked (negative selection). In the homology-enriched scores described in this work, correct decoys could be up-weighted by conserved interfaces in the homologous decoys and, at the same time, incorrect decoys could be discredited by statistically incompatible, clashing, or incomplete homologous decoys (since insertions in reference to the query structures were not modelled). We decided to first explore the simplest explanations, namely, deletions and/or clashes at the interface of homologs that would pull down the average score of the incorrect decoys. However, this does not seem to be the main driving force of ISC-h^10^’s success over ISC, as the number of gaps or the number of clashes (defined as heteroatom contacts under 1.5 Å) at the interface of homologous decoys do not strongly correlate with the given scores. Additionally, ranking using only the repulsive van der Waals component of the Rosetta score (fa_rep) performs very poorly in comparison to other scoring schemes with at most 34 out of 752 cases with correctly identified near-natives in the top 10 (supplementary Table 5-8). Finally, IES-h, SPP-h, or ISC-h variants where only the worst homologous decoys are taken into account when scoring each query decoy showed systematically worse performance than using the full range of homologous decoys for each query decoy (supplementary Table 5-8). This means that the performance of the homology-enriched scores is positively driven by the recognition of correct decoys rather than the exclusion of incorrect decoys through the presence of clashes or gaps.

Improvement of the SOAP-PP and Rosetta ISC scoring functions by homology enrichment is significant (supplementary Figure 5-2) and consistent over difficulty categories (supplementary Table 5-9). When splitting results over PPI4DOCK difficulty categories, we observe that the strongest relative gain for the SPP-h and ISC-h homology-enriched scores compared to their versions without homology occurs on “very_easy” cases, followed by “easy” cases (supplementary Table 5-9). A few cases are gained in the “hard” category, but the “very_hard” category remains largely inaccessible to the tested scores, even though our benchmark is limited to cases where at least one near-native decoy was sampled in the top 10,000 FRODOCK2.1 decoys (there are only 16 such “very_hard” cases). Consensus scoring also consistently improves results over the “very_easy”, “easy” and “hard” categories, in order of decreasing improvement. We hypothesise that correct ranking of very_easy and easy decoys is mainly dependent on the ability to score positively native-like models while more difficult categories would also require integration of flexibility, an ongoing challenge of protein docking (Desta, et al., 2020; Torchala, et al., 2013).

In this work, we developed a strategy to enrich scoring functions with evolutionary information by including atomic-level models for as few as ten homologs. This strategy improves the performance of several scores with different properties: InterEvScore (supplementary Table 5-10), SOAP-PP, and Rosetta ISC. This means that homology enrichment can in principle be applied to any scoring function with at most a ten-fold increase in runtime. This enrichment works with a very small number of sequences compared e.g. to the large MSAs needed by covariation methods to pick up coevolutionary signal, highlighting complementarity between the two approaches, which may be exploited by using additional DCA-derived constraints, e.g. in intermediate cases with a few hundred homologous sequences (Cong, et al., 2019; Simkovic, et al., 2017). The docking success boost also opens interesting perspectives regarding the large-scale application of structural prediction to interaction networks. Finally, with the rise of machine learning techniques in computational biology, one can expect interesting future developments using these approaches to further enhance the extraction of (co)evolutionary signal from coMSAs.

## 5 SUPPLEMENTARY INFORMATION

### 5.1 Supplementary methods

#### 5.1.1 Docking parameters

In the docking pipeline based on FRODOCK2.1, all parameters were set to default except for the following. Docking with the frodock executable used the “-t O” option for “other” complexes (not enzyme and not antibody-antigen). Clustering with frodockcluster was run with the −d 4 option, i.e. setting a LRMSD threshold of 4 Å for clustering.

#### 5.1.2 Alternative evaluation metrics: DockQ and nDCG

A more recent evaluation of decoy quality is given by the continuous DockQ score (Basu and Wallner, 2016), a metric going from 0 (bad quality) to 1 (high quality), which closely reflects the already-existing CAPRI quality categories. To integrate the DockQ score into a general performance measurement over our benchmark, we made use of the discounted cumulative gain (DCG) as in (Geng, et al., 2019). The DCG for each case is calculated as follows:

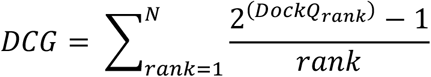

where rank is the rank of the decoy, DockQ_rank_ is the DockQ score of the decoy with that rank and N is the top N decoys that are taken into account for this measurement. The 1/rank factor gives more importance to the quality of the top scoring decoys. An ideal DCG (iDCG) is also calculated in order to normalise the DCG by reordering all decoys by decreasing DockQ score. A final normalised value (nDCG) for each case is deduced by dividing the DCG by the iDCG and can be extrapolated into a single value by calculating the average nDCG over all cases in the benchmark. Note that to speed up computations, decoys with a fraction of native contacts under 0.1 were given a default DockQ score of 0.

#### 5.1.3 Scoring functions

We employed an in house implementation of SOAP-PP that enables much more efficient scoring since decoy coordinates do not need to be explicitly generated. Note that only a slight reduction in performance on the 752 benchmark cases compared to the original SOAP-PP implementation has been observed (supplementary Table 5-11).

We also re-implemented InterEvScore for efficiency reasons. We introduced two variations compared to the best original InterEvScore (Andreani, et al., 2013): we defined interface contacts through distance thresholds (“distance mode”), instead of tessellation (“alpha mode” in InterEvScore) and we took evolutionary information into account for all interface residues instead of apolar patches only (so-called “standard mode” in the original implementation). InterEvScore outputs several scoring variants; here, we used the 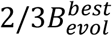 and the 2*B*^*best*^ (Andreani, et al., 2013). In 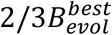, each interface residue contributes to the final score through the potential of its best 2- or 3-body contact and the potential of its equivalents in the homolog sequences. 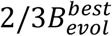 was found to perform best when scoring with homolog sequences (InterEvScore with implicit homology) (Andreani, et al., 2013) and thus was used in this context. 2*B*^*best*^ was used when scoring explicitly modelled side-chain models of our homologs (InterEvScore with explicit homology, IES-h). Indeed, we found that 3-body potentials are less discriminative than 2-body potentials in the context of explicitly modelled decoys (supplementary Table 5-12).

We use Rosetta 3.8 (version 2017.08.59291) and the beta_nov15 Rosetta score. Before scoring with Rosetta ISC, we perform high-resolution interface side-chain optimisation by using ‘use_input_sc’ and ‘docking_local_refine’ options of Rosetta’s docking_protocol executable. We also tried adding the ‘dock_min’ option (for even more conservative modelling and shorter scoring runtimes) but scoring results were degraded.

#### 5.1.4 Details on coMSA calculation

Compared to the original PPI4DOCK database (Yu and Guerois, 2016), coMSAs were slightly adjusted by realigning the first sequence (query) with all other sequences (considered as a block) using MAFFT (Katoh and Standley, 2013).

When building reduced coMSA^40^ from the readjusted PPI4DOCK coMSAs, coMSAs that already had under 40 sequences before the hhfilter step were not filtered.

The 10 sequences in coMSA^10^ were selected from coMSA^40^ as follows: Euclidian division was performed of the number of sequences in the coMSAs^40^ (including the query) over 10 with q and r, the quotient and remainder of this division. Starting from the first sequence, the next sequence is selected every q+1 for the first r steps, then every q until the end, including the last sequence resulting in 11 sequences with the first being the query and other 10, the homolog sequences.

#### 5.1.5 Threading models

The PPI4DOCK benchmark contains docking targets based on unbound homology models of pairs of binding partners for which an experimental complex structure is available. The use of homology modelling for unbound partners enables to expand the benchmark, by alleviating the need to identify complexes for which experimental structures of the interface and the exact two binding partners have been solved. This makes the benchmark larger, but as a counterpart, in PPI4DOCK the unbound structures used for docking are themselves, homology models.

In a docking context where we know the structures of the unbound partners, we would build homology models for all sequences in the coMSA by using the two query structures as modelling templates. However, since in PPI4DOCK the unbound query structures are themselves homology models, this would mean building a model by using a homology model as a template, and we felt this succession of modelling steps would lead to a loss in model precision. Therefore, the templates used for threading coMSA sequences were the unbound templates used to build the PPI4DOCK unbound models.

Template protein sequences were directly extracted from their structures and aligned onto the coMSAs using MAFFT (sequence-profile alignment) (Katoh and Standley, 2013) from which the pairwise homolog-template alignments were directly extracted. coMSAs were stripped down to positions that were covered by the query sequence. In order to ensure that the template structure exactly matched the template sequences in the stripped pairwise alignments, both template sequences were re-aligned using clustalw (Larkin, et al., 2007), and identified irrelevant residues in the template structure were removed.

Threading implies that the side-chains of our homologs are mapped very conservatively onto the query template structure.

### 5.2 Supplementary results

#### 5.2.1 Supplementary tables

**Table 5-1:**
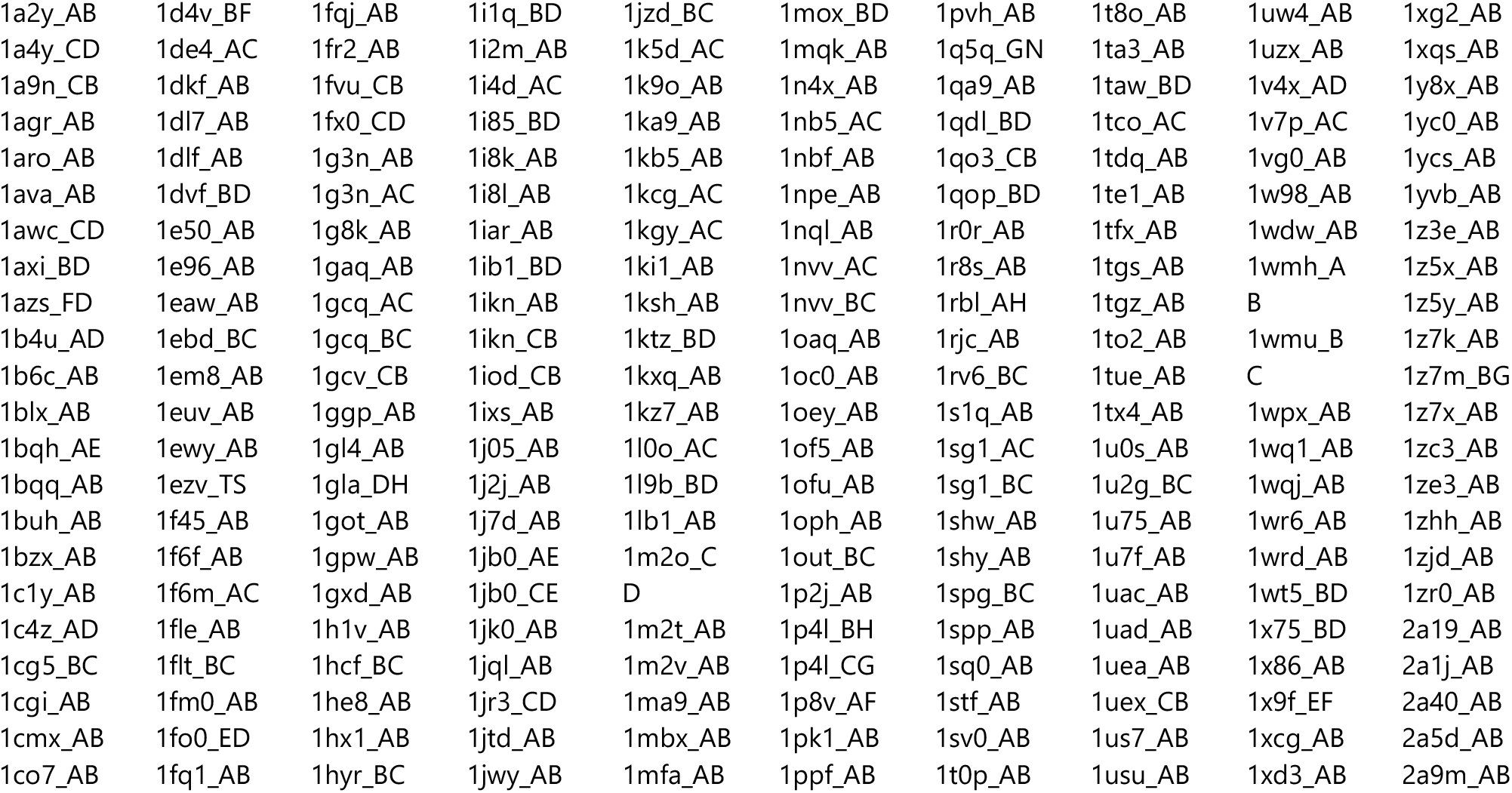
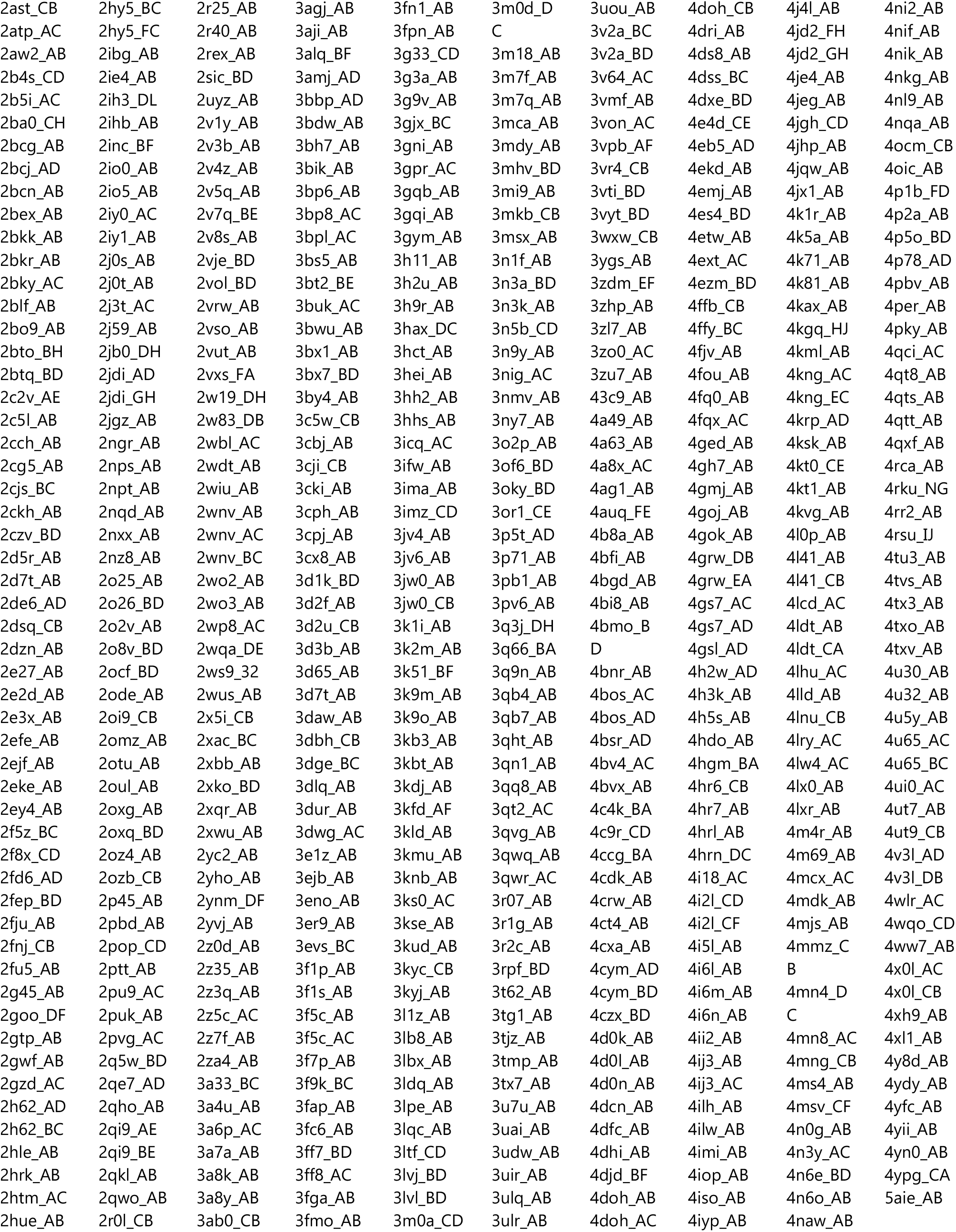
List of the 752 docking cases used as a benchmark set in this study. This subset of the 1417 cases in PPI4DOCK contains all cases with at least 10 sequences in the coMSAs and at least one acceptable decoy in the top 10,000 FRODOCK2.1 decoys.

**Table 5-2:**
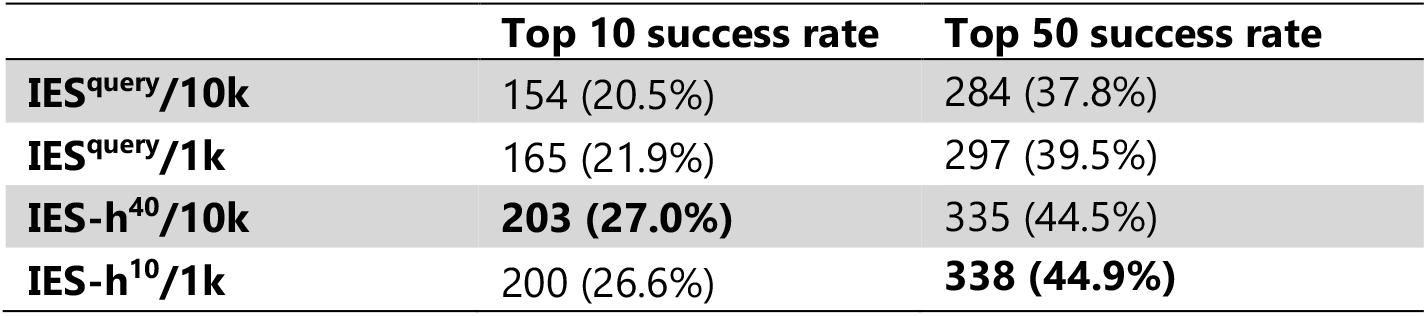
InterEvScore statistical potential. The IES^query^ score represents only the statistical potential part of InterEvScore (2B^best^) without any evolutionary information, used to rerank either the top 10,000 (10k) or the top 1,000 (1k) FRODOCK2.1 decoys. These results are shown for comparison with the homology-enriched IES-h variants described in the main results.

|  | Top 10 success rate | Top 50 success rate |
| --- | --- | --- |
| **IES<sup>query</sup>/10k** | 154 (20.5%) | 284 (37.8%) |
| **IES<sup>query</sup>/1k** | 165 (21.9%) | 297 (39.5%) |
| **IES-h<sup>40</sup>/10k** | **203 (27.0%)** | 335 (44.5%) |
| **IES-h<sup>10</sup>/1k** | 200 (26.6%) | **338 (44.9%)** |

**Table 5-3:** Scoring performance of homology-enriched SCORES. Scoring performance of ISC on query decoys only and using the threaded homology models (ISC-h^10^) on top 1,000 FRODOCK2.1 decoys (1k) and coMSA^10^ as well as the performance of SPP-h^40^ and IES-h^40^ on top 10,000 (10k) with coMSAs^40^ and the performance of SPP-h10 and IES-h10 on 1,000 FRODOCK2.1 decoys with coMSAs^10^ for easier comparison. Performances were measured as the top 10 success rate on 752 benchmark cases. This table is the same as Table 3-4 except that it includes coMSA^40^/10k success rates for comparison purposes.

|  | Top 10 success rate |  | Top 50 success rate |  |
| --- | --- | --- | --- | --- |
|  | coMSA<sup>40</sup>/10k | coMSA<sup>10</sup>/1k | coMSA<sup>40</sup>/10k | coMSA<sup>10</sup>/1k |
| **IES-h** | 203 (27.0%) | 200 (26.6%) | 335 (44.5%) | 338 (44.9%) |
| **SPP-h** | 228 (30.3%) | 227 (30.2%) | **365 (48.5%)** | **362 (48.1%)** |
| **ISC** | / | 157 (20.9%) | / | 301 (40.0%) |
| **ISC-h** | / | **259 (34.4%)** | / | **361 (48.0%)** |

**Table 5-4:** Numbers and timescales (on one CPU) of various elements and programmes. Times and numbers correspond to measurements on our 752-case PPI4DOCK benchmark. Decoys and docking mentioned below all refer to FRODOCK2.1 docking. The number of decoys generated per case changes according to the size of the complex, it averages at 9,651 with a maximum threshold of 10,000. Docking and decoy generation times are size-dependent but an average value is shown below.

|  |  |
| --- | --- |
| **Number of cases in our benchmark** | 752 |
| **Average number of sequences in our coMSAs** | 134 |
| **Average number of residues per case (receptor + ligand)** | 389 |
| **Maximum number of decoys generated in docking** | 10,000 |
| **Average number of decoys per case** | 9,651 |
| **Docking time with FRODOCK2.1** | 45 min - 1 h |
| **Structure generation time for 1,000 decoys with FRODOCK2.1** | 1 min |
| **Threading time with Rosetta per structure** | 1-2 min |
| **SOAP-PP scoring time for 1,000 decoys** | 1 min |
| **Original SOAP-PP scoring time for 1,000 decoys** | 15 min |
| **InterEvScore scoring time for 1,000 decoys** | 1 min |
| **Rosetta's ISC scoring time for 1,000 decoys** | 12 h 30 |
| **Consensus calculation time per case** | 20 s (3 scores) –<br>20 min (5 scores) |

**Table 5-5:** Top 1 and top 5 compared to top 10 success rates for consensus scores.

|  | Top 1 success rate | Top 5 success rate | Top 10 success rate |
| --- | --- | --- | --- |
| **Cons<sup>3</sup>** | 95 (12.6%) | 190 (25.3%) | 241 (32.0%) |
| **Cons<sup>3</sup>-h** | **113 (15.0%)** | 228 (30.3%) | 271 (36.0%) |
| **Cons<sup>4</sup>-h/150h** | 104 (13.8%) | 223 (29.7%) | 276 (36.7%) |
| **Cons<sup>4</sup>-h/1k** | 111 (14.8%) | 230 (30.6%) | 282 (37.5%) |
| **Cons<sup>5</sup>-h/150h** | 109 (14.5%) | 230 (30.6%) | 289 (38.4%) |
| **Cons<sup>5</sup>-h/1k** | **113 (15.0%)** | **247 (32.8%)** | **304 (40.4%)** |

**Table 5-6:** Performance with a more stringent near-native definition. Top 10 success rate with near-natives defined as being of at least Medium quality according to CAPRI criteria.

|  | Top 10 success rate | Top 50 success rate |
| --- | --- | --- |
| **FD** | 61 (8.1%) | 103 (13.7%) |
| **IES/10k** | 49 (6.5%) | 84 (11.2%) |
| **IES<sup>40</sup>/10k** | 50 (6.6%) | 87 (11.6%) |
| **IES-h<sup>40</sup>/10k** | 60 (8.0%) | 112 (14.9%) |
| **IES-h<sup>10</sup>/1k** | 66 (8.8%) | 107 (14.2%) |
| **SPP/10k** | 60 (8.0%) | 101 (13.4%) |
| **SPP-h<sup>40</sup>/10k** | 87 (11.6%) | 145 (19.3%) |
| **SPP-h<sup>10</sup>/1k** | 85 (11.3%) | 136 (18.1%) |
| **ISC/1k** | 50 (6.6%) | 93 (12.4%) |
| **ISC/150h** | 70 (9.3%) | 138 (18.4%) |
| **ISC-h<sup>10</sup>/1k** | 94 (12.5%) | 130 (17.3%) |
| **ISC-h<sup>10</sup>/150h** | **99 (13.2%)** | **159 (21.1%)** |
| **Cons<sup>3</sup>** | 62 (8.2%) | / |
| **Cons<sup>3</sup>-h** | 76 (10.1%) | / |
| **Cons<sup>4</sup>-h/150h** | 77 (10.2%) | / |
| **Cons<sup>4</sup>-h/1k** | 84 (11.2%) | / |
| **Cons<sup>5</sup>-h/150h** | 84 (11.2%) | / |
| **Cons<sup>5</sup>-h/1k** | 86 (11.4%) | / |

**Table 5-7:** Performance in terms of top 150 nDCG. Average nDCG were calculated and normalised over the top 150 decoys for each individual scores over 752 cases (see section 5.1.2).

|  | Top 150 success rate | nDCG /150 | nDCG /150<br>(excluding cases<br>with nDCG = 0) |
| --- | --- | --- | --- |
| **FD** | 387 | 0.118 | 0.147 |
| **IES/10k** | 377 | 0.135 | 0.180 |
| **IES<sup>40</sup>/10k** | 371 | 0.134 | 0.180 |
| **IES-h<sup>40</sup>/10k** | 417 | 0.157 | 0.195 |
| **IES-h<sup>10</sup>/1k** | 431 | 0.165 | 0.201 |
| **SPP/10k** | 444 | 0.138 | 0.157 |
| **SPP-h<sup>40</sup>/10k** | 455 | 0.180 | 0.207 |
| **SPP-h<sup>10</sup>/1k** | 458 | 0.186 | 0.213 |
| **ISC/1k** | 437 | 0.115 | 0.137 |
| **ISC/150h** | **476** | 0.149 | 0.169 |
| **ISC-h10/1k** | 451 | 0.182 | 0.213 |
| **ISC-h10/150h** | **476** | **0.208** | **0.236** |

**Table 5-8:** Performance of the repulsive term in Rosetta’s score and ISC-h^10^/1k on the worst third or worst homologs. Top 10 success rate of the fa_rep van der Waals repulsive terme in Rosetta’s scoring without (fa_rep /1k) and with homology through threaded homologs (fa_rep-h^10^/1k) as well as ISC-h^10^/1k using only the worst scoring third of homologs selected for each decoy individually (ISC-h^10/w3^/1k) or the worst scoring homolog for each decoy (ISC-h^10/w1^/1k) over 752 cases.

|  | Top 10<br>success rate |
| --- | --- |
| **fa_rep/1k** | 9 (1.2%) |
| **fa_rep-h<sup>10</sup>/1k** | 34 (4.5%) |
| **ISC/1k** | 157 (20.9%) |
| **ISC-h<sup>10</sup>/1k** | **259 (34.4%)** |
| **ISC-h<sup>10/w3</sup>/1k** | 227 (30.2%) |
| **ISC-h<sup>10/w1</sup>/1k** | 200 (26.6%) |
| **SPP/10k** | 183 (24.3%) |
| **SPP-h<sup>40</sup>/10k** | 228 (30.3%) |
| **SPP-h<sup>40/w3</sup>/10k** | 207 (27.5%) |
| **SPP-h<sup>40/w1</sup>/10k** | 188 (25.0%) |

**Table 5-9:**
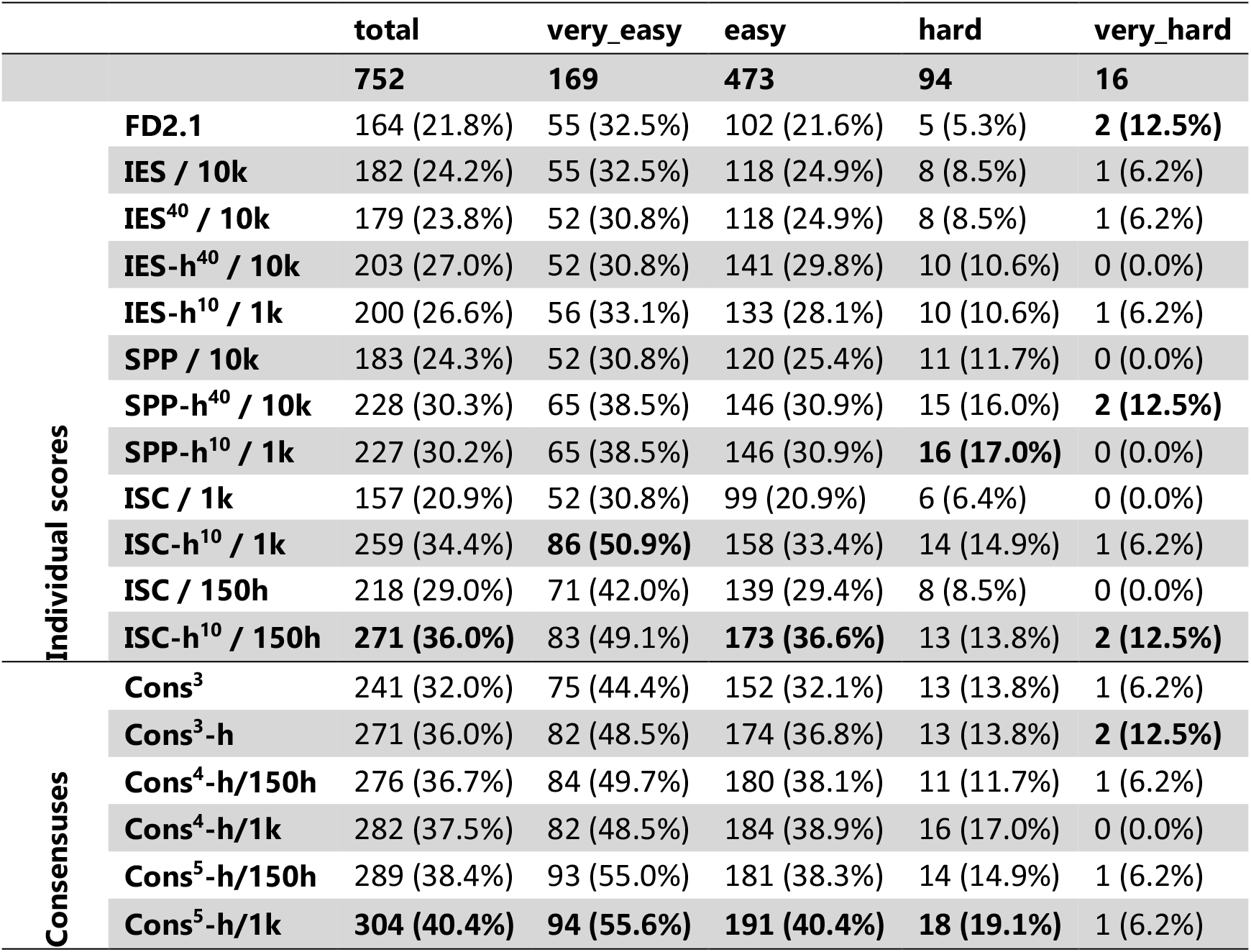
Performance over PPI4DOCK difficulty categories. Top 10 success rates separated over the four difficulty categories in our benchmark for FRODOCK2.1, InterEvScore and its threaded-homology variants, SOAP-PP and ISC and their evolutionary variants and the six consensus scores presented in section 3.5. Performances were measured on 752 benchmark cases.

**Table 5-10:** Performance of consensus scores including InterEvScore implicit homology scoring. Performance of three- and four-way consensus scores in terms of top 10 success rates on 752 benchmark cases. Scores used in Cons^3^ were SOAP-PP on the top 10,000 or top 1,000 FRODOCK2.1 decoys (SPP/10k or SPP/1k), InterEvScore on the top 10,000 or top 1,000 FRODOCK2.1 decoys (IES/10k or IES/1k) and FRODOCK2.1 (FD2.1). Scores used in Cons^4^ were SPP/10k, IES/10k, FRODOCK2.1 and Rosetta interface score on the top 1,000 FRODOCK2.1 decoys (ISC/1k). Performances of individual scores used in the consensuses are reported in terms of top 10 and top 50 success rates, since consensus calculation relies on the top 50 decoys ranked by each component score.

| Score | Top 10 success rate | Top 50 success rate |
| --- | --- | --- |
| FD2.1 | 164 (21.8%) | 292 (38.8%) |
| IES/10k | 182 (24.2%) | 287 (38.2%) |
| IES/1k | 196 (26.1%) | 295 (39.2%) |
| SPP/10k | 183 (24.3%) | **328 (43.6%)** |
| SPP/1k | 187 (24.9%) | 295 (39.2%) |
| Cons<sup>3</sup> | **241 (32.0%)** | / |
| ISC/1k | 157 (20.9%) | 301 (40.0%) |
| Cons<sup>4</sup> | 235 (31.2%) | / |

We try to improve the baseline consensus performance by incorporating Rosetta’s physics-based interface score (ISC) (section 2.2). As Rosetta scoring is more computationally expensive than the other two scores (about 750 times slower than SOAP-PP and InterEvScore calculations), we score only the top 1,000 decoys (as ranked by FRODOCK2.1) with ISC. This score is denoted ISC/1k as opposed to IES/10k and SPP/10k. As such, ISC is individually less well-performing than the other scores in terms of top 10 success rate, even when InterEvScore and SOAP-PP are computed only on the top 1,000 FRODOCK2.1 decoys (supplementary Table 5-10). However, the top 50 success rate is higher for ISC/1k than for any other individual score, except for SOAP-PP calculated on 10,000 decoys (supplementary Table 5-10). Despite this, integrating the top 50 decoys ranked by ISC/1k with the top 50 of the other three scores into a four-way consensus, denoted Cons^4^, slightly degrades performance compared to Cons^3^ (supplementary Table 5-10) while strongly increasing computation time.

**Table 5-11:** Performances as reported in the InterEvDock2 paper. Top 10 success rates of original scores in InterEvDock2 with percentages calculated over the same 752 cases compared with equivalent scores in this article. Original InterEvScore was run on the original PPI4DOCK coMSA and on the realigned coMSAs used throughout the present study (see section 5.1.4). Original SOAP-PP was run using the much slower Python implementation from the original publication.

|  | Top 10 success rate of original scores in InterEvDock2 | Top 10 success rate of new scores |
| --- | --- | --- |
| **FRODOCK2.1** | 164 (21.8%) | 164 (21.8%) |
| **InterEvScore** | 171 (22.7%) (original coMSAs)<br>177 (23.5%) (realigned coMSAs) | 182 (24.2%) |
| **SOAP-PP** | 194 (25.8%) | 183 (24.3%) |
| **3-way consensus** | 239 (31.8%) | 241 (32.0%) |

**Table 5-12:** Performances of InterEvScore with 2-body and 2/3-body potentials. Top 10 success rates of InterEvScore with complete coMSAs (IES) on 10,000 decoys, InterEvScore using homology models (IES-h) on coMSA^40^ and 10,000 decoys and on coMSA^10^ and 1,000 decoys using only 2-body potentials or 2- and 3-body potentials.

| | $2/3B^{best}$ | $2B^{best}$ |
| --- | --- | --- |
| **IES/10k** | 182 (24.2%) | 164 (21.8%) |
| **IES/1k** | 196 (26.1%) | 192 (25.5%) |
| **IES<sup>query</sup>/10k** | 147 (19.5%) | 154 (20.5%) |
| **IES<sup>query</sup>/1k** | 172 (22.9%) | 165 (21.9%) |
| **IES-h<sup>40</sup>/10k** | 161 (21.4%) | **203 (27.0%)** |
| **IES-h<sup>10</sup>/1k** | 182 (24.2%) | 200 (26.6%) |

### 5.2.2 Supplementary figures

**Figure 5-1:**
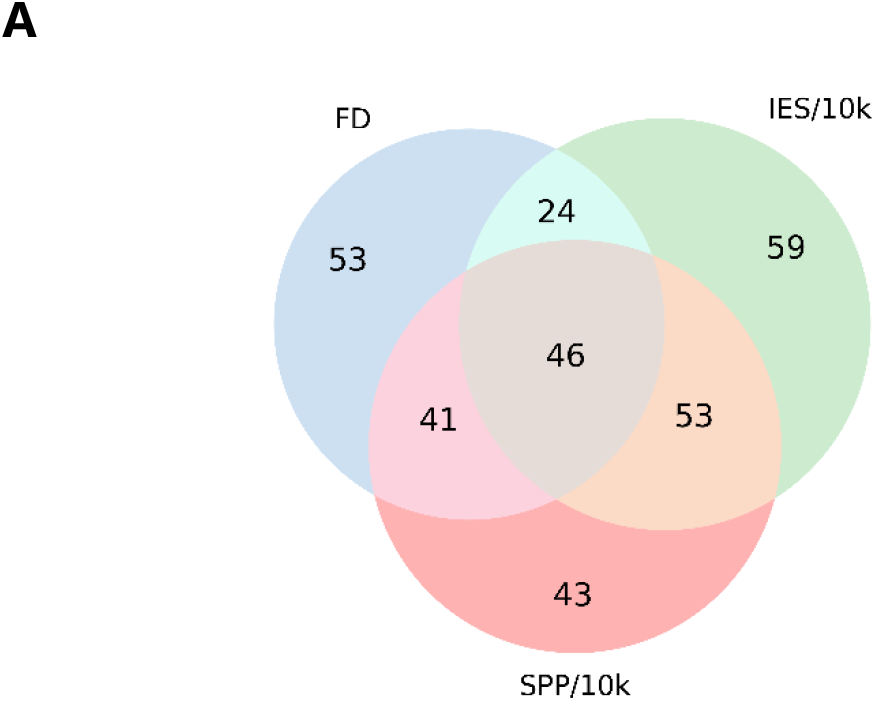
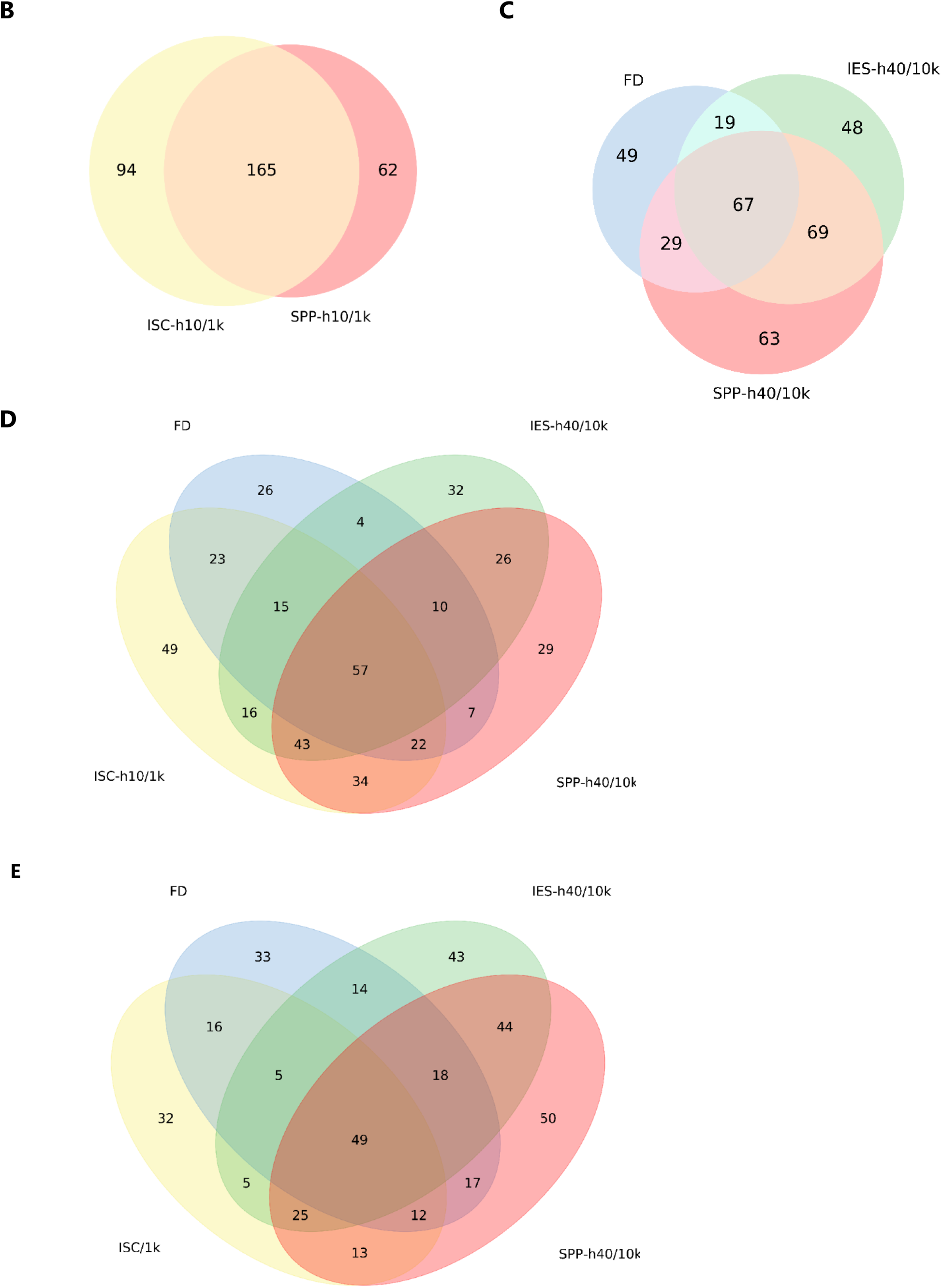
Venn diagrams between scores. Top 10 success rate intersections between scores on 752 cases. FD: FRODOCK2.1, IES: InterEvScore on complete coMSAs, SPP: SOAP-PP and ISC: Rosetta interface score. /10k and /1k denote that 10,000 and 1,000 decoys were scored. −h10 and −h40 denote homology-enriched scores with 10 or 40 homolog models (coMSA^10^ or coMSA^40^).

**Figure 5-2:**
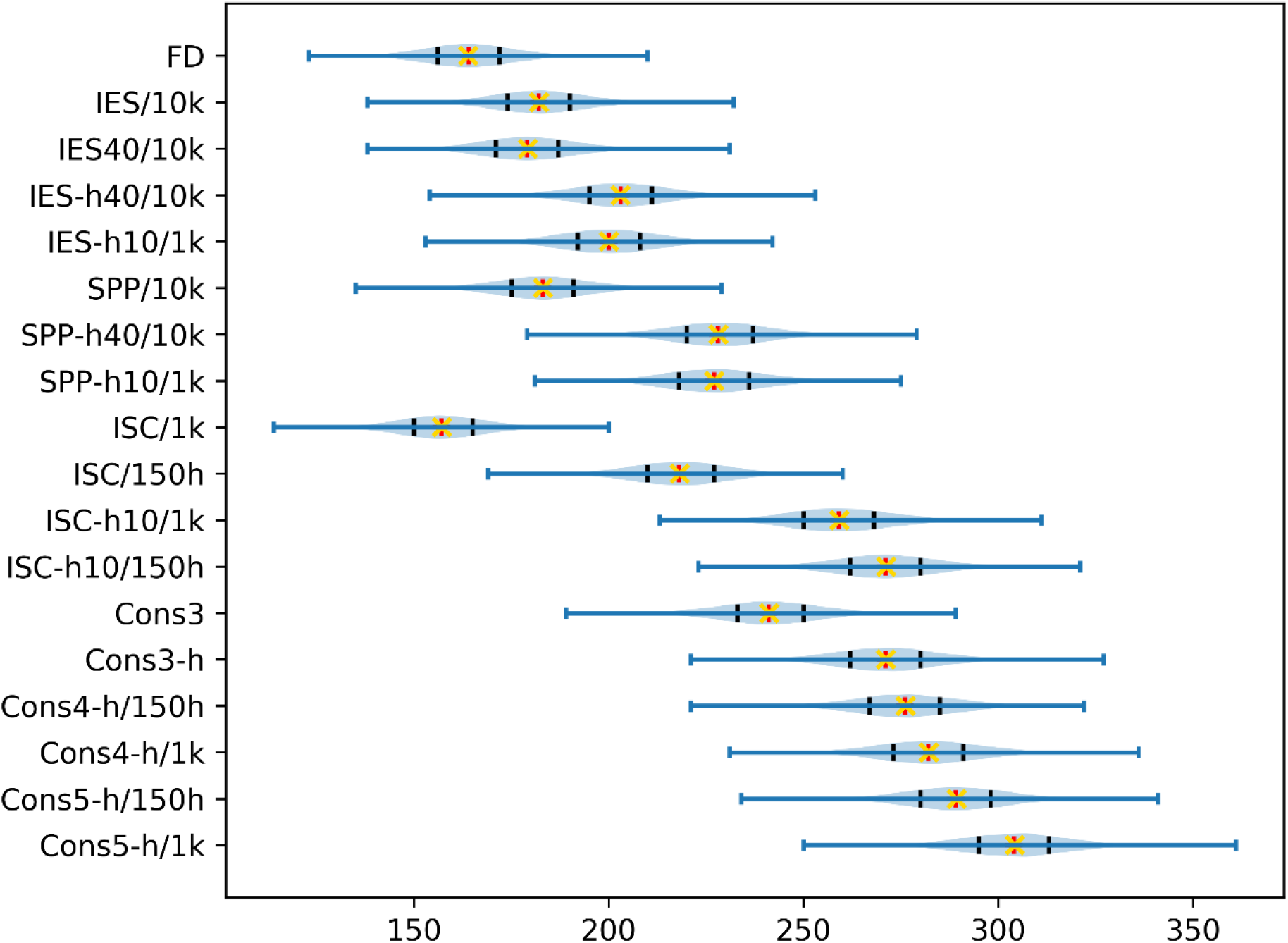
Bootstrap performance distributions. Bootstrap top 10 success rate distributions for 10,000 iterations over the 752 cases in our benchmark (blue). Measured top 10 success rates are marked in red and average success rates over all bootstrap iterations are marked as yellow crosses. Black bars indicate 25th and 75th percentiles of the bootstrap distribution. A two-sample t-test with unequal variances (Welch’s t-test) on all score pairs in this plot systematically outputs p-values < 10^−10^ except for Cons^3^-h against ISC-h^10^/150h, thus all distribution means are statistically different relative to each other except for these two scores.

## Acknowledgements

Benchmarking was done partly through granted access to the HPC resources of CCRT under the allocations 2018-7078 and 2019-7078 by GENCI (Grand Equipement National de Calcul Intensif). We thank Arnaud Martel for his help with setting up the data web page.

## Notes

**Funding** This work was supported by Agence Nationale de la Recherche [ANR‐15‐CE11‐0008 to R.G., ANR-18-CE45-0005 to J.A]; IDEX Paris-Saclay [IDI 2017 to C.Q.]; MINECO [BFU2016-76220-P to P.C.]; and AEI/FEDER, UE [PID2019-109041GB-C21 to P.C.].

http://biodev.cea.fr/interevol/interevdata/

### Competing Interest Statement

The authors have declared no competing interest.

